# Single-cell gene set enrichment analysis and transfer learning for functional annotation of scRNA-seq data

**DOI:** 10.1101/2022.10.24.513476

**Authors:** Simona Pellecchia, Gaetano Viscido, Melania Franchini, Gennaro Gambardella

## Abstract

Although an essential step, the functional annotation of cells often proves particularly challenging in the analysis of single-cell transcriptional data. Several methods have been developed to accomplish this task. However, in most cases, these rely on techniques initially developed for bulk RNA sequencing or simply make use of marker genes identified from cell clustering followed by supervised annotation. To overcome these limitations and automatise the process, we have developed two novel methods, the single-cell gene set enrichment analysis (scGSEA) and the single cell mapper (scMAP). scGSEA combines latent data representations and gene set enrichment scores to detect coordinated gene activity at single-cell resolution. scMAP uses transfer learning techniques to repurpose and contextualise new cells into a reference cell atlas. Using both simulated and real datasets, we show that scGSEA effectively recapitulates recurrent patterns of pathways’ activity shared by cells from different experimental conditions. At the same time, we show that scMAP can reliably map and contextualise new single cell profiles on a breast cancer atlas we recently released. Both tools are provided in an effective and straightforward workflow providing a framework to determine cell function and significantly improve annotation and interpretation of scRNA-seq data.

## INTRODUCTION

Single-cell RNA sequencing (scRNA-seq) delivers unprecedented opportunities for measuring gene expression at genome-wide scale and single-cell resolution. It provides a cost-effective way to study cellular tissue composition (1–5), dynamic processes during cell developmental stages (6), or the role of transcriptional heterogeneity in pathological conditions and in response to treatments (7). Indeed, scRNA-seq is becoming the leading technique for transcriptome profiling with large atlases of cells now routinely released from labs worldwide (1–8). Although scRNA-seq confers the advantage of measuring gene expression at the granular resolution of individual cells, the data produced with this sequencing technique are extremely noisy and zero-inflated (9). This property arises from both the low amount of mRNA available in a single cell and the limited capture efficiency of the technology. Thus, computational tools initially built to analyse bulk RNA-seq are often inadequate for scRNA-seq, and ad-hoc tools need to be developed to address the peculiarities of single-cell data. The significant increase in scRNA-seq studies raised several computational challenges, such as the development of automatic methods for functional annotation of scRNA-seq helpful for cell type assignment or disease diagnosis.

In the last decade, a plethora of tools have been developed to summarize regulated gene expression profiles into simplified functional categories useful for the annotation and interpretation of bulk RNA-seq data (10–16). Among these, the gene set enrichment analysis (GSEA) (11) is probably the most used. GSEA is a statistical tool that aims to measure coordinated activity of *a priori* defined gene sets (*i*.*e*., pathway) starting from a ranked list of genes usually obtained from differential expression (DE) analysis. By aggregating individual DE genes into pathways, GSEA projects DE results in a robust and easily interpretable biological space where additional analytics tools can be later applied. Currently, the application of GSEA to single cell data remains challenging (17) and only few methods are currently available, but are not efficient (18, 19). Therefore, there is an urgent need to develop novel computational methods capable of scoring the activity of a pathway within a reasonable time scale. On the other hand, the availability of large single-cell reference atlases comprising up to millions of cells across different species, conditions, tissues, or organs provides an unprecedented opportunity to use “re-purposing” techniques for the functional annotation of cells. Indeed, transfer learning (TL) techniques (20) are becoming increasing popular in the analysis of high-throughput -omics datasets including single-cell datasets (21–24). TL is a machine learning technique that uses joint embedding to integrate a query set of objects into a given reference and to contextualize them with the metadata associated with the reference elements.

Here, we present two novel ready-to-use pipelines tailored to scRNA-seq data that can be used to automatize cell functional annotation: scGSEA (single-cell gene set enrichment analysis) and scMAP (single-cell Mapper). scGSEA is a statistical framework for scoring coordinated gene activity in individual cells to automatically determine the pathways are active in a cell. scMAP is a TL algorithm to map a query set of cell transcriptional profiles on top of an existing reference atlas and contextualize the new data with the reference metadata. Both methods are based on non-negative matrix factorization (NMF) (25), a popular matrix decomposition method, that can be solved in a very computationally efficient manner (26). We validated scGSEA to identify pathway activity in both a simulated dataset and a real dataset comprising cells during various reprogramming stages (6) or drug treatment (27). We also tested the ability of scMAP in mapping novel sequenced cells from several conditions, including cells sequenced in different batches or another lab with a different sequencing technique. Both tools were developed in the framework of the gficf package (9, 28), an R package we recently developed for the normalization, visualization and clustering of single-cell data that takes advantage of text-mining approaches and available at https://github.com/gambalab/gficf.

## RESULTS

### Gifcf package overview

We recently introduced an R package named *gficf* useful for normalization of 3’ single-cell transcriptional data and the identification of biomarker genes across multiple experimental conditions or cell types (7, 9, 28). Our tool builds on a data transformation model named Gene Frequency – Inverse Cell Frequency (*i*.*e*., gf-icf) derived from the Term Frequency - Inverse Document Frequency (*i*.*e*., tf-idf) approach. TF-IDF is a statistical measure extensively used in the fields of text analysis and machine learning applied to Natural Language Processing (NLP) for quantifying the relevance of a word (*i*.*e*., gene) in a document (*i*.*e*., cell) amongst a comprehensive collection of documents (*i*.*e*., scRNA-seq dataset) (29, 30). When applying this model to scRNAseq data, the relevance of a gene increases proportionally to its expression in the cell but is offset by the frequency of the gene in the population of sequenced cells (28), so that only genes highly expressed in a small fraction of the cells are selected as the most relevant.

Briefly, the gf-icf pipeline (9, 28) starts from a set of single-cell transcriptional profiles and consists of the following steps: (*i*) cell quality control (QC) and filtering; (*ii*) rescaling of gene expression profiles of each cell to sum one (GF step) after raw count normalization (31); (*iii*) cross-cell normalization, to assign higher scores to rarely expressed genes than commonly expressed genes within each cell (ICF step); (*iv*) an L2 rescaling step to normalize gf-icf values; (*v*) linear dimensionality reduction of the data (*i*.*e*., PCA(32) or NMF(26)) to condense the complexity of the dataset into a lower-dimensional space (*vi*) non-linear dimensionality reduction (*i*.*e*., t-SNE (33) or UMAP (34)) of the data for its visualization and finally (*vii*) several downstream analyses including cell clustering and differential expression (9) (Figure 1A). GF-ICF method is implemented as an open-source R package, freely available at https://github.com/gambalab/gficf.

**Figure 1.**
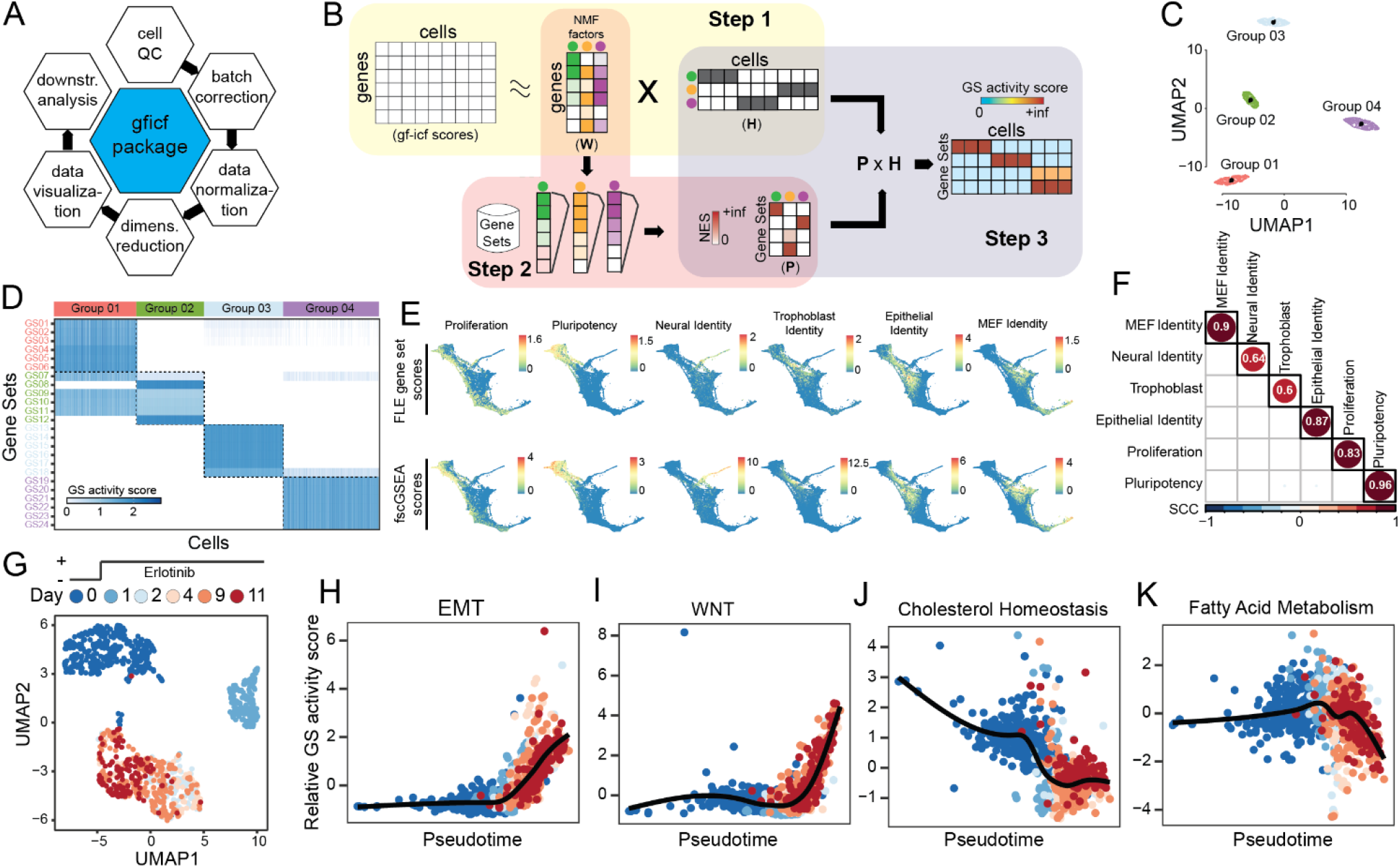
Single cell gene set enrichment analysis overview and performances. (**A**) GFICF package overview. (**B**) Single cell gene set enrichment analysis pipeline. (**C**) UMAP plot of 5,000 simulated cells grouped in four distinct groups. (**D**) Reconstructed activity of 24 simulated pathways across the 5,000 cells in (C). In the heatmap pathways are along rows while simulated cells along columns. Cells are ordered according to their group of origin. (**E**) Comparison between scGSEA pathway scores and signature scores originally computed by Schiebinger et al. on 251,203 single-cell profiles collected during differentiation stages. First row shows original gene set scores computed by Schiebinger et al. using wot phyton package. Second row shows gene set scores computed with scGSEA tool in the gficf R package. Each column represents a different gene set. Scores were plotted on the original FLE (force-directed layout) coordinates published by Schiebinger et al. (**F**) Spearman Correlation Coefficient (SCC) between scGSEA scores and wot package signature scores across the 251,203 single-cell transcriptional profiles in (E). (**G**) UMAP representation of 1,044 cells subject to eleven days of consecutive erlotinib treatment. Cells are color-coded according to sequenced day (i.e., 0, 1, 2, 4, 9, and 11 days). Single cell transcriptional profiles were normalized with gficf package. (**H**) EMT activity scores against inferred cell pseudo-time using the activity scores of 50 hallmark gene sets downloaded from MSigDB. Cells are color-coded as in (G). (**I-K**) Same as (H) but for wnt, cholesterol and fatty acid pathways respectively.

Here we updated the *gf-icf* package and implemented two novel functionalities: scGSEA and scMap.

### scGSEA for the reconstruction of pathway activity at single cell resolution

We aimed to construct a bioinformatic method that could measure the activity of an *a priori* defined collection of gene sets (*i*.*e*., pathways) at the single cell resolution. To this end, we developed a robust and fast single-cell Gene Set Enrichment Analysis (scGSEA) algorithm that takes advantage of the informative biological signals spreading across the latent factors of gene expression values obtained from non-negative matrix factorization (Methods) (25, 26). The scGSEA method starts from a set of single-cell expression profiles and a collection of gene sets and scores their cumulative expression (*i*.*e*., pathway activity) in each of the profiled cells and it is divided into three main steps (Figure 1B). In the first step, we use NMF to decompose GF-ICF normalized data into two positive related matrices containing gene weights (**W**) and cell weights (**H**). Each column of **W** or row of **H** defines a factor representing complex biological processes that recur throughout the set of sequenced cells (35, 36). While, at the same time, all features define the latent space amongst genes and cells. In the second step, we infer which biological processes each latent factor is associated with by performing GSEA (37) against each column of the **W** matrix. Since the values in each column of the **W** matrix are continuous weights describing the relative contribution of a gene in each inferred factor, a positive and significant value of the enrichment score (ES) for a specific pathway implies the factor is related to it. At the end of this step, the **W** matrix is transformed into a novel positive matrix **P** describing the pathways’ contribution in the form of normalized enrichment scores (NES) for each latent factor. Finally, in the third step, we infer the pathway’s activity level in each profiled cell by multiplying the **P** matrix reconstructed in step two with the **H** matrix obtained using the NMF method in step one. The rationale behind this last step is that each column of the matrix **H** describes the relative contribution of a cell across the factors. Thus, a cell with a high weight for a specific factor is assumed to share the phenotype or biological process related to that factor (36). Consequently, the activity level of a pathway in a cell can be computed as the weighted sum of the normalized pathway’s enrichment scores across the factors shared by the cell.

We performed simulations to assess the effectiveness of scGSEA in recapitulating the activity level of a pathway at single cell level (see methods for details). By using the splatter package (38) we generated a zero-inflated count dataset composed of 5,000 cells grouped in four distinct populations (Figure 1C) through a gamma-Poisson distribution using parameters inferred from real dataset. Then, we assigned 24 overlapping gene sets of different sizes (Supplementary Table 01) to be exclusively expressed in each cluster (*i*.*e*., six specific gene sets per cluster). As shown in Figure 1D, we found that scGSEA can identify in each group of cells the activity of the 6 specific simulated pathways with some of these correctly predicted active also in other groups of cells due to their partial overlap with other genes sets (Supplementary Figure 01).

To demonstrate the reliability of the scGSEA method on a real dataset, we applied it to the cell reprogramming scRNA-seq data in Schiebinger et al. (6). This dataset comprises 251,203 single-cell transcriptional profiles collected at half-day intervals across 18 days of reprogramming by ectopic expression of OKSM (a.k.a. Oct4, Klf4, Sox2, and Myc) transcription factors. In this work, the authors constructed seven curated gene signatures to score cells as pluripotent-, epithelial-, trophoblast-, neural-, MEF-like and proliferative. Hence, we applied our scGSEA method to this dataset with these seven gene sets as the input. As shown in Figure 1E-F, we found a high degree of correlation (avg. 0.8) between the predicted scGSEA pathway’s activity level and the pathway expression computed by Schiebinger et al. (6).

Next, we investigated whether we could use scGSEA scores to infer cell trajectories and reconstruct dynamics of the key pathways driving resistance to EGFR inhibitors in non-small-cell lung carcinoma (NSCLC). Several studies have suggested that acquired resistance mechanisms to EGFR inhibitors involve the compensatory activation of redundant signalling pathways that share effectors or downstream modulators of the EGFR signalling cascade, thus bypassing EGFR inhibition. Different pathways can serve as alternative routes for reactivation of signalling downstream of inhibited EGFR, including MET, IGF-1R, PI3K-AKT-mTOR, BRAF/RAS and Wnt signalling pathways (39–42), all able to sustain cell survival, proliferation, migration, and epithelialmesenchymal transition (EMT).

Therefore, we downloaded the scRNA-seq dataset published by Aissa et al. (27) comprising 1,044 PC9 lung cells (Figure 1G) that were subject to eleven days of consecutive erlotinib treatment and sequenced at six different time points (i.e., 0, 1, 2, 4, 9, and 11 days). We then applied scGSEA to score the activity of 50 MSigDB hallmark pathways (43) across the cells. The resulting scores were used as input to reconstruct the dynamic activity of these pathway by applying a pseudotime algorithm (44) (Supplementary Figures 02). As shown in Figure 1H,I we found strong upregulation of EMT and Wnt b-catenin signalling pathways in erlotinib tolerant cells, while erlotinib tolerant cells showed a potent inhibition of genes related to cholesterol and fatty acid metabolism, as also reported by Aissa et al. (Figure 1J,K).

These results show how scGSEA could recapitulate recurrent patterns of pathways’ activity shared by hundreds of thousands of cells from multiple conditions and during dynamic processes like cell reprogramming or drug treatments.

### Mapping a single cell transcriptional profile on a reference atlas

We aimed to construct a computational method that starting from the transcriptional state of a cell, could be integrated, and contextualized within a larger reference cell atlas by using the metadata associated to the reference cells. To this end, we developed scMAP (single-cell Mapper), a transfer learning algorithm that combines text mining data transformation and a k-nearest neighbours’ (KNN) classifier (methods) to map a query set of single-cell transcriptional profiles on top of a reference atlas. Our strategy consists of three main steps, as schematised in Supplementary Figure 03: (i) we first normalize the query cell profiles with the GF-ICF method by using the ICF weights learned by the reference atlas; (ii) we then project normalized cell profiles to the NMF (or PC) sub-space of the reference atlas before mapping them onto its UMAP embedding space; (iii) finally, we use the KNN algorithm to contextualize mapped cells.

To test scMAP, we took advantage of the single-cell atlas of breast cancer we have recently released (7). This atlas comprises 35,276 individual cells from 32 breast cancer cell lines covering all four major breast tumour subtypes (*i*.*e*., LuminalA, LuminalB, Her2-positive and Basal Like). Hence, we first applied GF-ICF tool (28) on these cells to generate a reference UMAP embedding space from the either from the top 100 NMF factors (Figure 2A) or the top 50 PC (Figure 2B). Next, we used three approaches to test the accuracy of our method in correctly mapping the very cells in the breast cancer atlas. First, we used a cross-validation approach where we randomly divided the 35,276 single-cell transcriptional profiles in different proportions of training and test cells (*i*.*e*., from 10 to 90% of the cells in each cell line). With this approach, we use training cells to reconstruct the reference atlas and the remaining cells to measure the performances of the remapping algorithm. Second, we re-sequenced with the DROP-seq platform 1,683 cells of two cell lines included in the atlas and mapped the transcriptome of these cells onto it. Third, we tested our mapping strategy on 16,683 single cell transcriptional profiles from 11 cell lines included in the atlas but sequenced by other laboratories with 10X genomics platform (8, 45). In all cases the performances of the mapping algorithm were tested by using either NMF or PCA as cell sub-space before remapping them into the UMAP embedding space. After remapping, the label of a cell was assigned using the closest 101 cells (7) (see methods).

**Figure 2.**
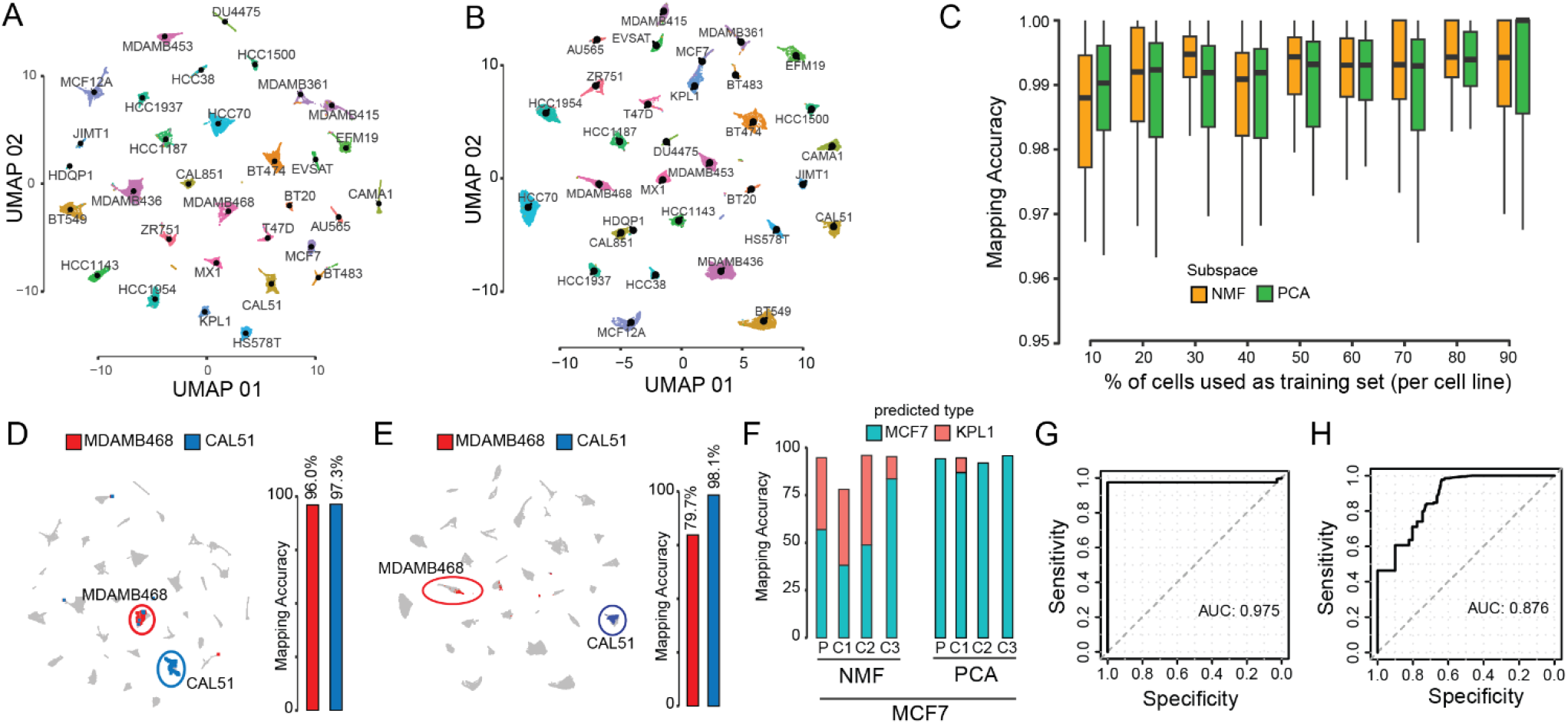
Single cell mapping accuracy evaluation. (**A**) UMAP representation of 35,276 cells from 32 breast cancer cell-lines using as cell subspace NMF. Cells are color-coded according to their cell-line of origin. Single cell transcriptional profiles were normalized with gficf package. (**B**) Same as (A) but using PC as cell sub-space (**C**) Accuracy evaluation of mapping method using cross-validation approach using as cell sub-space either NMF (orange) or PCA (green). Each boxplot display accuracy distribution in classifying the 32 cell lines but using as a training set the percentage of cells indicated on the x-axis. Accuracy is defined as the number of correctly classified cells over the total number of mapped cells. (**D**) Left plot; mapping of the MDAMB468 and CAL51 cells after they were re-sequenced with drop-SEQ technology. Right plot; accuracy of the mapping method in classifying re-sequenced MDAMB468 and CAL51 cells. NMF cell subspace is used. (**E**) Same as (**D**) but using PC sub-space. (**F**) Accuracy evaluation of the mapping method on 14,372 single cell transcriptomes sequenced with 10x Chromium method for MCF7 parental (P) cell line and three derived subclones (C1,C2,C3) using as cell sub-space either NMF (left) or PCA (right). (**G**) Performance of the mapping method in classifying 2,311 single cell transcriptomes sequenced using the 10x Chromium method from eleven distinct breast cell lines cell. Performances are reported in terms ROC curve and AUC is also displayed. (**H**) Same as (G) but using PC cell sub-space.

As shown in Figure 2C, the median cross-validation approach’s accuracy of the mapping algorithm was 97% when using NFM as cell sub-space and 99% when using PC as sub-space. However, with the cross-validation approach we always use cells sequenced in the same batch. Thus, to avoid this possible bias, we re-sequenced 306 and 1,377 cells of MDAMB468 and CAL51 cell lines, respectively (see methods). As shown in Figure 2D, 96% of MDAMB468 and 97.3% for CAL51 cells were recognized and labelled correctly when using NFM as cell sub-space. While when using PC as subspace mapping accuracy was of 79.7% and 98.1% for MDAM468 and CAL51 respectively (Figure 2E).

Next, we investigated whether the sequencing technology could affect the efficacy of the mapping algorithm. Thus, we started mapping into the reference breast cancer cell-line atlas 14,372 single cell transcriptomes sequenced using the 10x Chromium method from the MCF7 cell line and three subclones derived from it (45). Considering that KPL1 has been documented as an MCF7 derivative cell line (46, 47), we found that 13,699 of the 14,372 cells (95.3%) were recognized correctly (Figure 2F) when using NFM as subspace. Using PC instead of NFM as a cell sub-space, the total accuracy was of 94.2% (Figure 2F). Finally, we mapped 2,311 single cell transcriptomes sequenced using the 10x Chromium method from eleven additional breast cell lines included in our atlas and published by Kinker et al. (8). Figure 2G shows classification performance on these cells in terms of ROC (Receiver operating characteristic) curve with an AUC of 0.975. The AUC of the ROC curve was 0.876 when the PC subspace was used instead of NFM (Figure 2H).

Overall, these analyses shows that the mapping approach and transfer learning strategy we developed provide reliable results with good accuracy.

## DISCUSSION

Single-cell RNA-seq provides a cost-effective way to study cell composition of tissues (1–5), cell developmental stages (6), and to elucidate the role of transcriptional heterogeneity in pathological conditions or in response to drug treatments (7). Indeed, large atlases of cells with their relative associated metadata are now routinely released. However, computational methods for the reconstruction of pathway activity at single cell level useful for their annotation or to contextualize novel sequenced cells in an already available and annotated atlas of cells remain challenging.

Finding an effective way to capture coordinated gene activity at the single-cell level is crucial for single-cell data transformation and subsequent analyses. Such transformation allows representation of cellular state in terms of activity levels of biological processes (*i*.*e*., set of genes) rather than through the expression levels of individual genes. Thus, allowing the projection of single-cell data to a quickly biologically interpretable space in which analytic approaches can later be applied. For example, such representations could be used to detect within the same cell type shifts in the portion of cells exhibiting an altered biological process between two phenotypes of interest. Thus improving the identification of dysregulated signalling pathways across pathological conditions, otherwise identified from differential expression (31, 48–52) or co-expression (19, 53–65) analyses that are still challenging for single cell datasets due to their zero-inflated nature (66–72).

Here, we introduced scGSEA, a novel efficient ready-to-use method that provides an effective and simple workflow for the measurement of pathway activity at single cell level. Our scGSEA takes advantage of the informative biological signals spreading across the colinearly optimized and additive collection of factors of gene expression values obtained from an NMF model. Indeed, when applied on bulk transcriptional datasets, NMF factors have already been shown to better capture patterns of coordinated gene activities compared to other matrix decomposition methods like SVD ranks or PCA components (36) Recently, NMF has been shown to greatly improve scRNA-seq data clustering and visualization thanks to its inbuilt ability to impute missing values and decompose data into additive parts (26). In applications, we demonstrated that our scGSEA has high accuracy in capturing coordinated gene activity at the single-cell level in both simulated and real dataset comprising hundreds of thousands of cells. However, scGSEA is a tool that leverages NMF expression latent factors to infer pathway activity at a single cell level. Thus, by design, it inherits both benefits and limitations of the NMF model. For example, a limit of this model, like other matrix decomposition techniques, is the choice of the exact number of factors to use. This choice can be made by either using a subjective approach like finding an inflection point in a curve of ranks against an objective (i.e., elbow plot) or with higher precision, but computationally demanding approaches, like jackstraw or k-fold cross-validation. However, when choosing the number of factors to use, to avoid information loss, it is advisable to adopt a cautious approach by selecting a higher number. Thus, we generally recommend using at least 100 NMF factors on a large dataset comprising more than 10,000 cells, otherwise 50 should suffice.

In this study we also presented scMAP, a transfer learning algorithm that combines text mining data transformation and the k-nearest neighbours’ algorithm to map a query set of cell transcriptional profiles on top of an existing cell atlas. Finding an effective way to “re-purpose” sequenced cells in an already annotated dataset of cells could be helpful in the study of different diseases. For example, we have already shown that we can use breast cancer cell lines for automatic cancer subtype classification starting from the single cell transcriptomic dataset of patient biopsies (7). Recently, it has also been shown that with transfer learning, we can use the knowledge of sensitive drugs for each cell line to predict the patient’s treatment once the patient’s cells were confidently mapped on the reference atlas (73). Here, we demonstrated our method has high accuracy in cell mapping even when we profile cells’ transcriptomes after several culture passages in a different batch or with a different sequencing technique. The proposed mapping method may be applicable in several scenarios; however, it is best suited when the query cells consist of cell types and experimental protocols close to the reference data. Finally, the number of shared genes between the query and reference cells can also impact mapping accuracy. We recommend using all available genes in the reference-building step to guarantee more extensive feature overlap between reference and query cells, which naturally increases the mapping quality.

In summary, we have updated our gficf R package with two novel functionalities, both useful for cell functional annotation. One allows to capture coordinated gene activity at the single-cell level, and another that can re-purpose newly sequenced cells into an already annotated reference dataset. Both tools are provided in an effective and straightforward workflow and are implemented in the framework of our open-source R package gficf available at https://github.com/gambalab/gficf.

## METHODS

### Cell culture

MDAMB468 and CAL51 cell lines used in this study were obtained from commercial providers and cultured in ATCC recommended complete media at 37°C and 5% CO2.

### scRNA library preparation, sequencing and alignment

Single cell transcriptomic of the MDAMB468 and CAL51 cell lines were performed with DROP-seq technology (74) and library preparation as described in Gambardella et al. (7). scRNA libraries were sequenced with NovaSeq 6000 machine using an SP 100 cycles flow cell. Raw reads pre-processing was performed using Drop-seq tools v2.3.0 and following the Dropseq Core Computational Protocol reported at http://mccarrolllab.org/dropseq. Briefly, raw reads were first filtered to remove all read pairs with at least one base in their cell barcode or UMI with a quality score less than 10. Then read 2 was trimmed at the 5’ end to remove any TSO adapter sequence, and at the 3’ end to remove polyA tails. Filtered reads were then fed to STARsolo tool v2.7.10a (https://github.com/cellgeni/STARsolo) to perform alignment, UMI deduplication and gene expression quantification. Hg38 human genome (primary assembly v40) downloaded from GENCODE (75) was used as a reference genome for read alignment. Only high depth cells with at least 2,500 UMI were retained and used to test our cell mapping tool. Alignment pipeline can be found at https://github.com/gambalab/dropseq.

### Single Cell Gene Set Enrichment Analysis (scGSEA)

To perform scGSEA, raw count matrix was first normalized with gficf package (28). Then, NMF was used to decompose gficf scores into two positive related matrices containing gene weights (**W**) and cell weights (**H**) where each column of **W** or row of **H** defines a latent actor *f*_*i*_. Non-negative matrix factorization was performed by using the fast parallel implementation can be found in RcppML R package (26) available at https://github.com/zdebruine/RcppML. Then gene set enrichment analysis was performed against each column of the **W** matrix using as input a pre-defined list of gene sets *S =* {*s*_1_, *s*_2_ *… s*_*n*_}. GSEA was performed using the R package fgsea (37) available at https://github.com/ctlab/fgsea. At the end of this process, a novel matrix **P** with the same number of columns (*i*.*e*., factors) of **W** and with the number or rows equal to the number of inputs used gene set is obtained. Each element of **P**_*i,j*_ contains the normalized enrichment scores of the pathway *s*_*i*_ related to the factor *f*_*j*_. Next, only positives and significant normalized enrichment scores with an FDR<0.05 were retained while all the other elements of **P** are put to zero. Finally, since elements of **P** describe the pathways’ contribution in each latent factor, the pathway’s activity level in each profiled cell is computed as the weighted sum of the normalized pathway’s enrichment scores across the factors shared by the cell. This corresponds simply to the dot product of the **P** and **H** matrices. The described pipeline is implemented in the function scGSEA of the gficf package.

### Simulated scRNA-seq profiles and gene sets

Splatter R package (38) with default parameters was used to generate a zero-inflated count dataset composed of 5,000 cells and 1,000 genes with cells grouped in four distinct populations. Next, six gene sets were simulated to be exclusively expressed in each group of simulated cells (*i*.*e*., 6 gene sets x 4 groups of cells = 24 simulated gene sets). The 24 gene sets comprised 2,432 unique genes that we added to the simulated dataset generated with splatter. A zero-inflate Poisson distribution with success probability equal to 50% and lambada value of 10 was used to simulate the expression of a gene set in a specific set of cells. This to simulate a moderate dropout and a relatively low gene count expression for each gene set.

### Pseudo-time analisys

scGSEA method was run using as imput the 1,044 PC9 single cell transcriptional profiles and the 50 hallmarks gene sets (v2022.1) downloaded from MSigDB (www.gsea-msigdb.org/gsea/msigdb). The obtained pathways’ activity matrix was used to run psupertime function of psupertime R package and infer the pseudo-time order of cells. Psupertime function was run with default parameters and using as labels their sequencing day (i.e., 0, 1, 2, 4, 9 or 11)

### Single Cell Mapper (scMAP)

UMAP embedding space of reference single cell breast cancer atlas was built from scratch with gficf package (28). Then, new sequenced cells were mapped on the reference breast cancer atlas following the strategy depicted in Supplementary Figure 02. Briefly, scRNA-seq profiles are first normalized with *gficf* strategy but using the ICF weight learned on the reference atlas and then projected to the existing NMF (or PC) sub-space using gene loadings learned from the breast cancer atlas. These values are then used as input of the *umap_transform* function of *uwot* package, which uses the UMAP reference model to map the new cells into the reference UMAP space. Finally, the cell line of origin associated with each mapped cell was predicted by using k nearest-neighbour algorithm from KernelKnn package (https://mlampros.github.io/KernelKnn). For all assignments k parameter was set to 101. Single cell mapping is implemented in the function scMAP of the gficf package.

### Public single-cell transcriptional dataset

The raw counts of the 35,276 single cell transcriptional profiles of the 32 breast cancer cell (7) used in this study were downloaded from figshare (https://doi.org/10.6084/m9.figshare.15022698). The 251,203 single-cell transcriptional profile from pluripotent stem cells (6) were obtained from GEO database with accession number GSE122662. The single cell transcriptional profiles of MCF7 cells and derived clones (45) were obtained from GEO database with accession number GSE114462. The 2,311 single cell the transcriptional profile across the eleven breast cancer cell lines (8) were obtained from GEO database with accession number GSE157220. PC9 lung cells were obtained from GEO database with accession number GSE149383 but only 1,044 cells with a total number of UMI greaten then 2,500 were used in this work.

## CODE AVAILABILITY

R package of gf-icf pipeline and examples of use are available at the following address https://github.com/gambalab/gficf. While scripts to reproduce main figures and analyses are available at the following address https://github.com/gambalab/scGSA_scMAP_manuscript.

## DATA AVAILABILITY

Single-cell RNA-seq of MDAMB468 and CAL51 cell lines can be found on the GEO database with accession number GSE214827.

## AUTHOR CONTRIBUTIONS

GG wrote the manuscript, conceived, and developed the tools, SP and GV performed NGS experiments while MF performed bioinformatics analysis.

