## Supplementary Figures for "Single-cell gene set enrichment analysis and transfer learning for functional annotation of scRNA-seq data"

Cell functional annotation with text mining approaches.

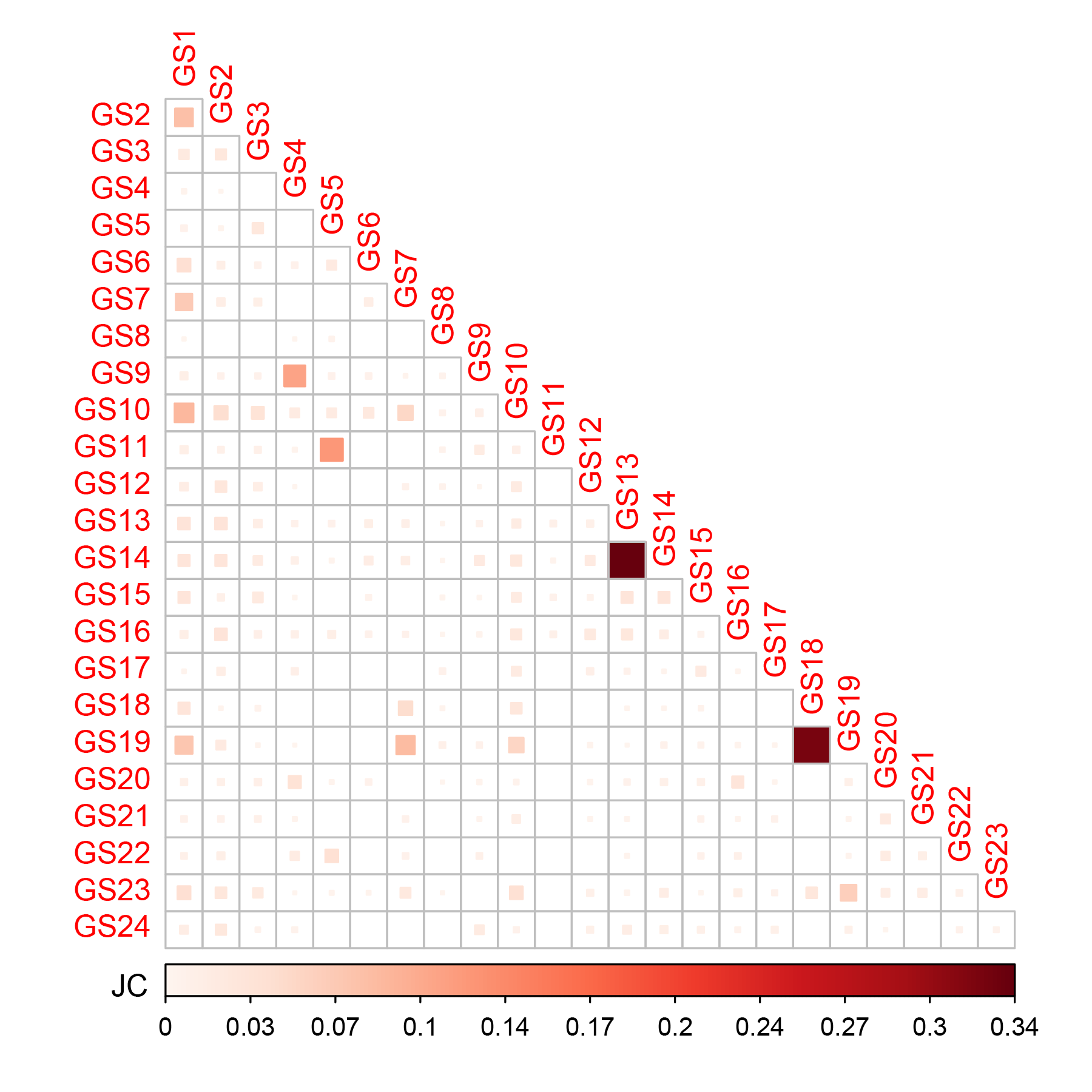

**Supplementary Figure 1 – Simulated gene sets overlap.** Jaccard Coefficient (JC) between the 24 simulated gene sets.

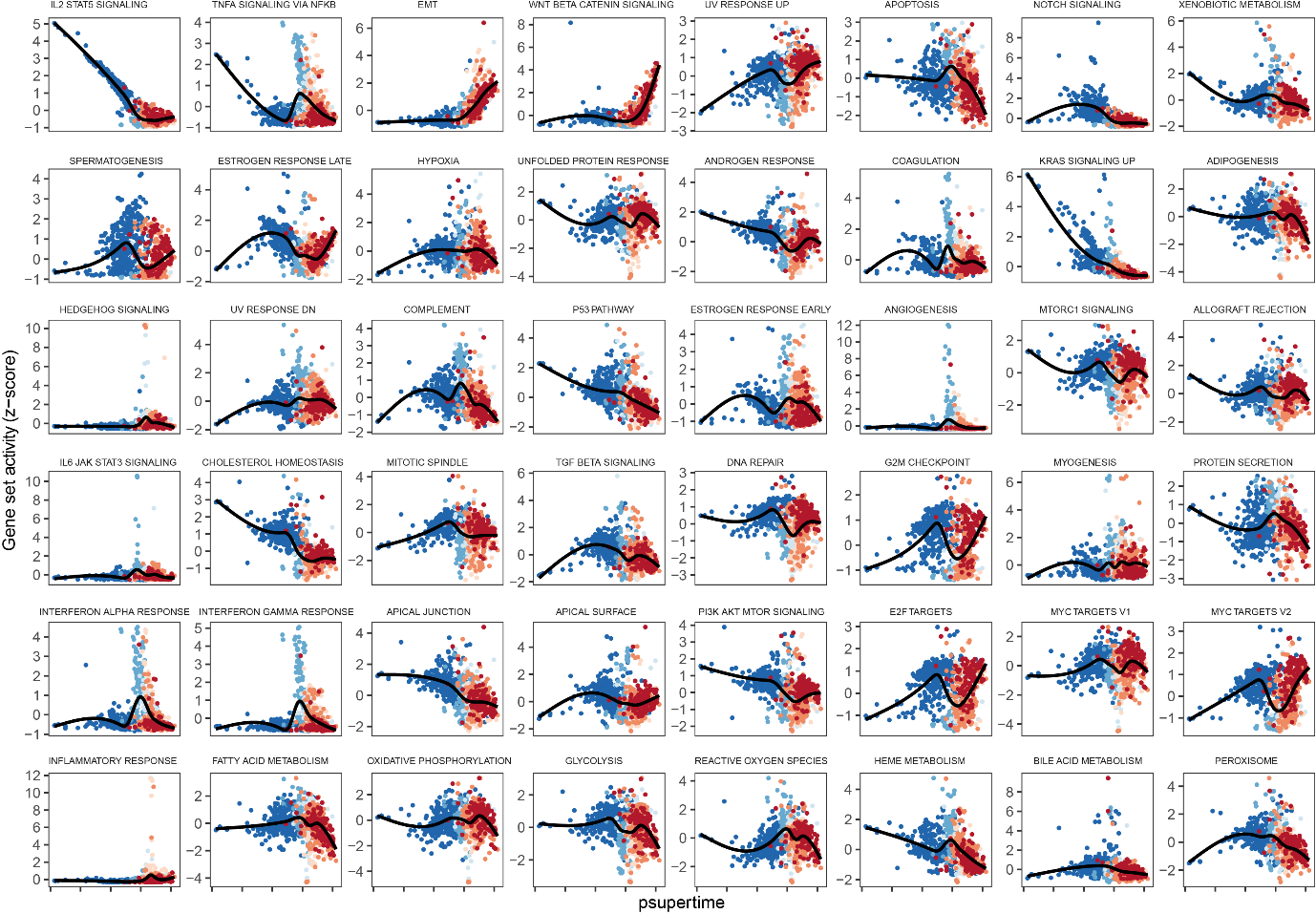

**Supplementary Figure 2 – Pathway activity scores over time.** Activity scores against inferred cell pseudo-time of the 50 hallmark gene sets from MSigDB.

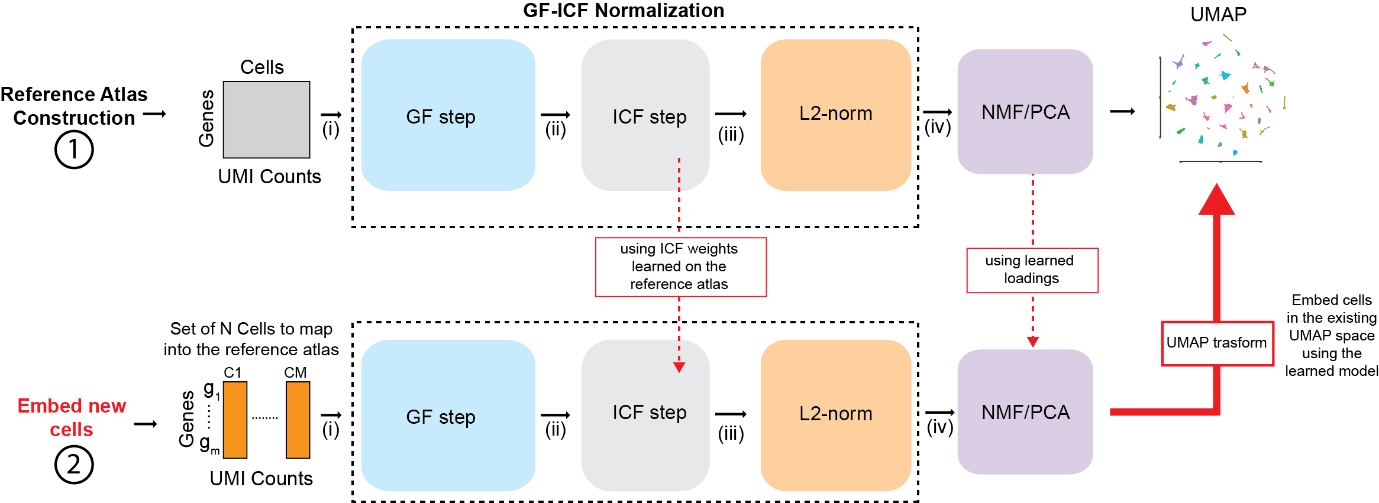

**Supplementary Figure 3 - Description of the single-cell mapping algorithm**. Following reference atlas construction (step 1), additional scRNA-seq profiles can be added to the atlas (step 2) by first normalizing scRNA-seq data with the gficf package using the ICF weights estimated during the atlas construction and then by projecting the normalised scRNA-seq to the NMF/PC space using gene loadings from the reference atlas. Finally, the umap_transform function of uwot package is used to embed the new cells into the reference UMAP space.
